# Detecting Differential Variable microRNAs via Model-Based Clustering

**DOI:** 10.1101/296947

**Authors:** Xuan Li, Yuejiao Fu, Xiaogang Wang, Dawn L. DeMeo, Kelan Tantisira, Scott Weiss, Weiliang Qiu

**Affiliations:** Department of Mathematics and Statistics, York University, Toronto, ON, Canada; Channing Division of Network Medicine, Brigham and Women’s Hospital, Harvard Medical School, Boston, USA

**Keywords:** F test, equal variance, EM algorithm, epigenetics, hepatocellular carcinoma

## Abstract

Identifying genomic probes (e.g., DNA methylation marks) is becoming a new approach to detect novel genomic risk factors for complex human diseases. The F test is the standard equal-variance test in Statistics. For high-throughput genomic data, the probe-wise F test has been successfully used to detect biologically relevant DNA methylation marks that have different variances between two groups of subjects (e.g., cases vs. controls). In addition to DNA methylation, microRNA is another mechanism of epigenetics. However, to the best of our knowledge, no studies have identified differentially variable (DV) microRNAs. In this article, we proposed a novel model-based clustering to improve the power of the probe-wise F test to detect DV microRNAs. We imposed special structures on covariance matrices for each cluster of microRNAs based on the prior information about the relationship between variance in cases and variance in controls and about the independence among cases and controls. To the best of our knowledge, the proposed method is the first clustering algorithm that aims to detect DV genomic probes. Simulation studies showed that the proposed method outperformed the probe-wise F test and had certain robustness to the violation of the normality assumption. Based on two real datasets about human hepatocellular carcinoma (HCC), we identified 7 DV-only microRNAs (hsa-miR-1826, hsa-miR-191, hsa-miR-194-star, hsa-miR-222, hsa-miR-502-3p, hsa-miR-93, and hsa-miR-99b) using the proposed method, one (hsa-miR-1826) of which has not yet been reported to relate to HCC in the literature.

## INTRODUCTION

Investigating the relationship between genomics and complex human diseases has greatly improved our understanding of the molecular mechanisms of, and the interplay of environmental factors and genomic factors to, the complex human diseases. High-throughput data from cutting-edge technologies have substantially facilitated the unbiased discovery of the genetic risk factors for many diseases. The standard approach to identify disease-associated genomic probes is to test if the mean expression (e.g., DNA methylation) between cases and controls is significantly different. In Statistics, variance is another important measurement, in addition to mean. The larger the variance is, the more information the data could provide. However, the information about variance has not been directly used to detect disease-associated genomic probes until recent years.

Several groups of researchers have recently identified DNA methylation marks that have different variances between cases and controls[1-3]. They observed that (1) for differentially variable DNA methylation marks the variability in cases is usually higher than that in controls; and (2) differentially variable DNA methylation marks are biologically relevant. DNA methylation is a type of epigenetics, which studies the heritable changes in organisms caused by regulating gene expression without changing genetic code. DNA methylation inhibits gene expression by adding a methyl group to the cytosine or adenine DNA nucleotides. Another type of epigenetics is microRNAs that are short noncoding 18-25 nucleotide long RNA and negatively regulate mRNA translation [4, 5]. However, to the best of our knowledge, no studies have investigated differential variability for microRNAs. The main objective of this article is to develop statistical methods to detect microRNAs differentially variable between cases and controls.

The F test is the classical method to test for equal variance between two groups of subjects, which evaluates whether the ratio of sample variances between two groups is significantly different from one. For high-throughput genomic data, such as DNA methylation data, the probe-wise F test could be used. That is, we first perform the F test for each probe to test for equal variance between cases and controls. We then calculate FDR-adjusted p-value to control for multiple testing, where FDR stands for false discovery rate. If the FDR-adjusted p-value < 0.05 for a DNA methylation mark, we then claim that this DNA methylation mark is differentially variable between cases and controls. The advantages of this probe-wise approach include flexibility (one model per probe) and easy-implementation. However, differentially variable probes might be governed by the same underlying mechanism. Statistically speaking, the variances of differentially variable probes might follow the same distribution. Similarly, the variances of non-differentially variable probes might also follow the same distribution. We hypothesize that these underlying distributions of variances could help us improve the power of the F test to detect differentially variable probes.

In this article, we propose a mixture of 3-component multivariate normal distributions to fit the expression levels of microRNAs to identify microRNAs differentially variable (i.e., having different variances) between cases and controls.

## METHOD

### Model

We assume that microRNAs belong to one and only one of the following three clusters:

(1) microRNAs having higher variances in cases than in controls (denoted the cluster as OV), (2) microRNAs having equal variances between cases and controls (denoted the cluster as EV), and (3) microRNAs having smaller variances in cases (denoted the cluster as UV). We follow Qiu et al. (2008)[6] to directly model the marginal distributions of microRNAs in the 3 clusters. In this article, we modified Qiu et al.’s (2008) marginal model[6] to allow the detection of DV probes. We assume that (1) data have been normalized to remove the effects of confounding factors, such as chip effect, and batch effect, etc., and (2) data have been transformed so that the distributions of microRNA expressions are close to normal distributions.

For a given microRNA, we denote *X*_*i*_ as the pre-processed expression for the *i*-th subject, *i*=1, …, *m*, where *m*=*m*_*c*_ + *m*_*n*_, *m*_*c*_ is the number of cases and *m*_*n*_ is the number of controls. For the *k*-th cluster (*k*=1, 2, or 3), we assume that the expressions of the *m*_*c*_ cases are identically distributed with mean *μ*_*kc*_ and variance 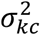. We assume that the expressions of the *m*_*n*_ controls are identically distributed with mean *μ*_*kc*_ and variance 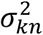. According to Qiu et al. (2008)[6], *X*_*i*_’s are marginally correlated with correlation *ρ*_*kc*_ for cases and *ρ*_*kn*_ for controls. We also assume that (1) cases and controls are independent, and (2) the *m*×1 random vector (*X*_1_,…, *X*_*m*_)^*T*^ follows a multivariate normal distribution. For the OV cluster, we require that 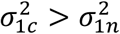. For UV cluster, we require that 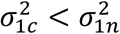. For the EV cluster, we require that 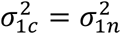. We allow the means and correlations are different between cases and controls in the EV cluster.

We used the EM algorithm[7] to estimate the model parameters μ_*kc*_, 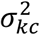, μ_*kn*_, and 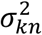. The posterior probability 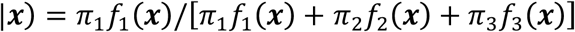 is used to assign the *g*-th microRNA to one of the 3 clusters, where *f*_*k*_(***x***) is the density function of the multivariate normal distribution for the *k*-th cluster. If *p*_*g1*_ is the largest posterior probability among *p*_*g*1_,*p*_*g*2_, and *p*_*g*3_, then the *g*-th microRNA will be assigned to the 1^st^ cluster (i.e., OV cluster). The Supplementary Document gives the details about the model and the corresponding parameter estimation procedure.

### Real datasets

We downloaded two microRNA datasets from NIH’s Gene Expression Omnibus (GEO)[8]: GSE67138 (https://www.ncbi.nlm.nih.gov/geo/query/acc.cgi?acc=GSE67138) and GSE67139 (https://www.ncbi.nlm.nih.gov/geo/query/acc.cgi?acc=GSE67139). Both datasets are from the same project that aims to detect microRNAs differentially expressed between human hepatocellular carcinoma (HCC) tumor tissues with and without vascular invasion. GSE67138 is the first batch containing 57 samples (34 invasive tumor tissues and 23 non-invasive tumor tissues), while GSE67139 is the second batch containing 120 samples (60 invasive tumor tissues and 60 non-invasive tumor tissues). The expression levels of microRNAs in both GEO datasets were measured by using Affymetrix Multispecies miRNA-1 Array (GPL8786). Both datasets contain 847 microRNAs.

We checked the data quality by visualizing the plot (Fig. A1) of percentiles across arrays and the plot (Fig. A2) of the first and second principal components. Both plots indicate the two datasets have been cleaned and have good quality (i.e., no apparent outlying microRNAs, outlying arrays, or technical batch effects). Hence, we directly use the two datasets in the further analyses. We regarded GSE67138 as the discovery set and GSE67139 as the validation set.

### Simulation

We conducted 4 sets of simulation studies. In the first set (denoted it as SimI), we generated microRNA data from the proposed marginal mixture model, where estimated model parameters for GSE67138 (i.e., the discovery set) are used as the true values of the model parameters (*π*_1_ = 0.10, *π*_2_ = 0.84, *π*_3_ = 0.06, *μ*_1*c*_ = −0.80, 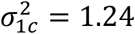, *ρ*_1*c*_ = −0.02, *μ*_1*n*_ = 0.55, 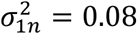, *ρ*_1*n*_ = 0.14). We generated 100 datasets, each of which has 1,000 microRNAs for 50 cases and 50 controls. Ten percent (100) of the 1,000 microRNAs are in the OV cluster. Six percent (60) microRNAs are in the UV cluster. The remaining 84% (840) microRNAs are in the EV cluster.

In the second set (denoted it as SimII), we generated microRNA data from a mixture of 3 multivariate t distribution with the same mean vectors and covariance matrices as those in SimI and with degrees of freedom 3. SimII is used to evaluate the performance of the proposed method when the normality assumption is violated.

In the third set (denoted it as SimIII) of the simulation studies, we generated microRNA data from the same model as that in SimI, except that the marginal correlations within subject-groups were set to zero (*ρ*_*kc*_ = 0 and *ρ*_*kn*_ = 0). SimIII is used to evaluate the performance of the proposed method when there are no marginal correlations.

In the fourth set (denoted it as SimIV) of the simulation studies, we generated microRNA data from the same model as that in SimII, except that the marginal correlations within subject-groups were set to zero (*ρ*_*kc*_ = 0 and *ρ*_*kn*_ = 0). SimIV is used to evaluate the performance of the proposed method when there are no marginal correlations and when the normality assumption is violated.

### Statistical Analysis

We compared the proposed method (denoted as gs) with ten existing equal-variance tests by using both the real datasets and the simulated datasets. The ten equal variance tests are: (1) F test (denoted as F); (2) Ahn and Wang’s score test[9] (denoted as AW); (3) Phipson and Oshlack’s AD test[10] (denoted as PO.AD); (4) Phipson and Oshlack’s SQ test[10] (denoted as PO.SQ); (5) Levene’s test[11] (denoted as L); (6) Brown and Forsythe’s test[12] (denoted as BF); (7) trimmed-mean-based Levene’s test[12] (denoted as Ltrim); (8) improved AW test based on Levene’s test[13] (denoted as iL); (9) improved AW test based on BF test[13] (denoted as iBF); and (10) improved AW test based on trimmed-mean-based Levene’s test[13] (denoted as iTrim). Except for the F test, the other 9 tests are robust to the violation of the normality assumption.

We first applied the 11 methods (the gs method and the 10 existing methods) to the discovery set (GSE67138) to detect microRNAs differentially variable between invasive tumors and non-invasive tumors. For the 10 existing methods, we obtained FDR-adjusted p-values. If a microRNA has FDR-adjusted p-value < 0.05, we claim that this microRNA has significantly different variances between invasive tumors and non-invasive tumors. We then applied the same procedure to the validation set (GSE67139). We claim that a microRNA is a validated DV microRNA (1) if the microRNA is DV in both discovery and validation sets, and (2) if the sign of the difference 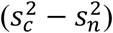 is the same in both datasets, where 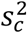 and 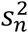 are sample variances for cases and controls, respectively. We next calculated the proportion of the validated DV microRNAs *pValid*=*n*_*12*_/*n*_*1*_, where *n*_*1*_ is the number of DV microRNAs in the discovery set (GSE67138) and *n*_*12*_ is the number of DV microRNAs in both the discovery set (GSE67138) and the validation set (GSE67139).

For the validated DV microRNAs detected by the gs method, we also check if they are validated differentially expressed (DE) microRNAs by using R Bioconductor package *limma*[14]. A microRNA is a validated DE microRNA if the FDR-adjusted p-value for testing equal mean expression between cases and controls is < 0.05 in both the discovery and validation sets and if the sign of the mean difference 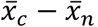 is the same in both discovery and validation sets, where 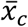 and 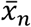 are the sample means of the cases and controls, respectively. Denote *S*_*DVonly*_ as the set of microRNAs that are validated DV, but not validated DE. Denote *S*_*DEonly*_ as the set of microRNAs that are validated DE, but not validated DV. Denote *S*_*both*_ as the set of microRNAs that are both validated DE and validated DV.

We applied miRSystem[15] to predict the target genes of microRNAs in each of the 3 sets: *S*_*DVonly*_, *S*_*DEonly*_, and *S*_*both*_. miRSystem also provides the enriched KEGG pathways for the predicted target genes.

For simulated datasets, we calculated the magnitude of agreement between the true cluster memberships of microRNAs and the detected cluster memberships by each of the 11 methods by using the Jaccard index[6, 16]. The maximum value of the Jaccard index is one, indicating perfect agreement. The minimum value of the Jaccard index is zero, indicating that the agreement is by chance.

## Results

For the real data analyses, the numbers of the DV microRNAs in the discovery set, and the numbers and proportions of the validated DV microRNAs are shown in Table 1. The gs method detected 132 DV probes based on the discovery set, 67 of which are validated in the validation set. Among the 67 validated DV microRNAs, 66 microRNAs are OV and only one microRNA is UV. The proportion of the validated DV microRNAs is 0.51 for the gs method, which is the highest among the 11 methods. For all the 11 methods, the number (nValid.OV) of the validated OV microRNAs is much larger than the number (nValid.UV) of the validated UV microRNAs. This observation is consistent with what observed by other researchers using DNA methylation data[3].

**Table 1:** Information about the validated DV microRNAs

| method | nSig | n.OV | n.UV | nValid | nValid.OV | nValid.UV | pValid |
| --- | --- | --- | --- | --- | --- | --- | --- |
| gs | 132 | 91 | 41 | 67 | 66 | 1 | 0.51 |
| F | 239 | 183 | 56 | 106 | 96 | 10 | 0.44 |
| AW | 55 | 46 | 9 | 26 | 26 | 0 | 0.47 |
| PO.AD | 148 | 113 | 35 | 59 | 54 | 5 | 0.40 |
| PO.SQ | 58 | 48 | 10 | 29 | 29 | 0 | 0.50 |
| L | 139 | 106 | 33 | 58 | 54 | 4 | 0.42 |
| BF | 53 | 37 | 16 | 24 | 24 | 0 | 0.45 |
| Ltrim | 79 | 57 | 22 | 34 | 33 | 1 | 0.43 |
| iL | 131 | 99 | 32 | 55 | 51 | 4 | 0.42 |
| iBF | 49 | 35 | 14 | 23 | 23 | 0 | 0.47 |
| iTrim | 72 | 52 | 20 | 31 | 30 | 1 | 0.43 |

nSig: the number of the DV microRNAs detected in the discovery set; n.OV: the number of OV microRNAs detected in the discovery set; n.UV: the number of UV microRNAs detected in the discovery set; nValid: the number of validated DV microRNAs; nValid.OV: the number of validated OV microRNAs; nValid.UV: the number of validated UV microRNAs; pValid = nValid/nSig.

We got 324 DE microRNAs based on the discovery set, among which 217 DE microRNAs were validated. There are only 7 microRNAs in *S*_*DVonly*_ (hsa-miR-1826, hsa-miR-191, hsa-miR-194-star, hsa-miR-222, hsa-miR-502-3p, hsa-miR-93, and hsa-miR-99b), the parallel boxplots of which are shown in Fig. A3. *S*_*DEonly*_ contains 157 microRNAs (Table A1), the parallel boxplots of which are shown in Fig. A4. *S*_*both*_ contains 60 microRNAs (Table A2), the parallel boxplots of which are shown in Fig. A5.

Based on the miRSystem analysis, there are 1,639 genes (Table A3) targeted by the 7 microRNAs in *S*_*DVonly*_, 8,141 targeted genes (Table A4) for the 157 microRNAs in *S*_*DEonly*_, and 6,893 targeted genes (Table A5) for the 60 microRNAs in *S*_*both*_.

The 1,639 genes targeted by the 7 microRNAs in *S*_*DVonly*_ are significantly enriched (RAW_P_VALUE < 0.05) in 6 KEGG pathways (CALCIUM SIGNALING PATHWAY, SALIVARY SECRETION, AMYOTROPHIC LATERAL SCLEROSIS (ALS), MAPK SIGNALING PATHWAY, PPAR SIGNALING PATHWAY, and ALZHEIMER’S DISEASE) (Table A6). The 8,141 genes targeted by the 157 microRNAs in *S*_*DEonly*_ are significantly enriched in only one KEGG pathway (ANTIGEN PROCESSING AND PRESENTATION) with RAW_P_VALUE=2.70E-2 (Table A7). The 6,893 genes targeted by the 60 microRNAs in *S*_*both*_ are enriched in two KEGG pathways (O-GLYCAN BIOSYNTHESIS and GLYCINE SERINE AND THREONINE METABOLISM) (Table A8).

For the first simulation study (SimI), almost all of the 100 values of the Jaccard index by the gs method are equal to one (the perfect agreement). Figure 1 showed that the gs method had significantly higher values of the Jaccard index than the 10 existing equal-variance tests. The pattern of the Jaccard index values is similar for SimIII (Figure 3), where subjects are marginally independent.

**Figure 1.**
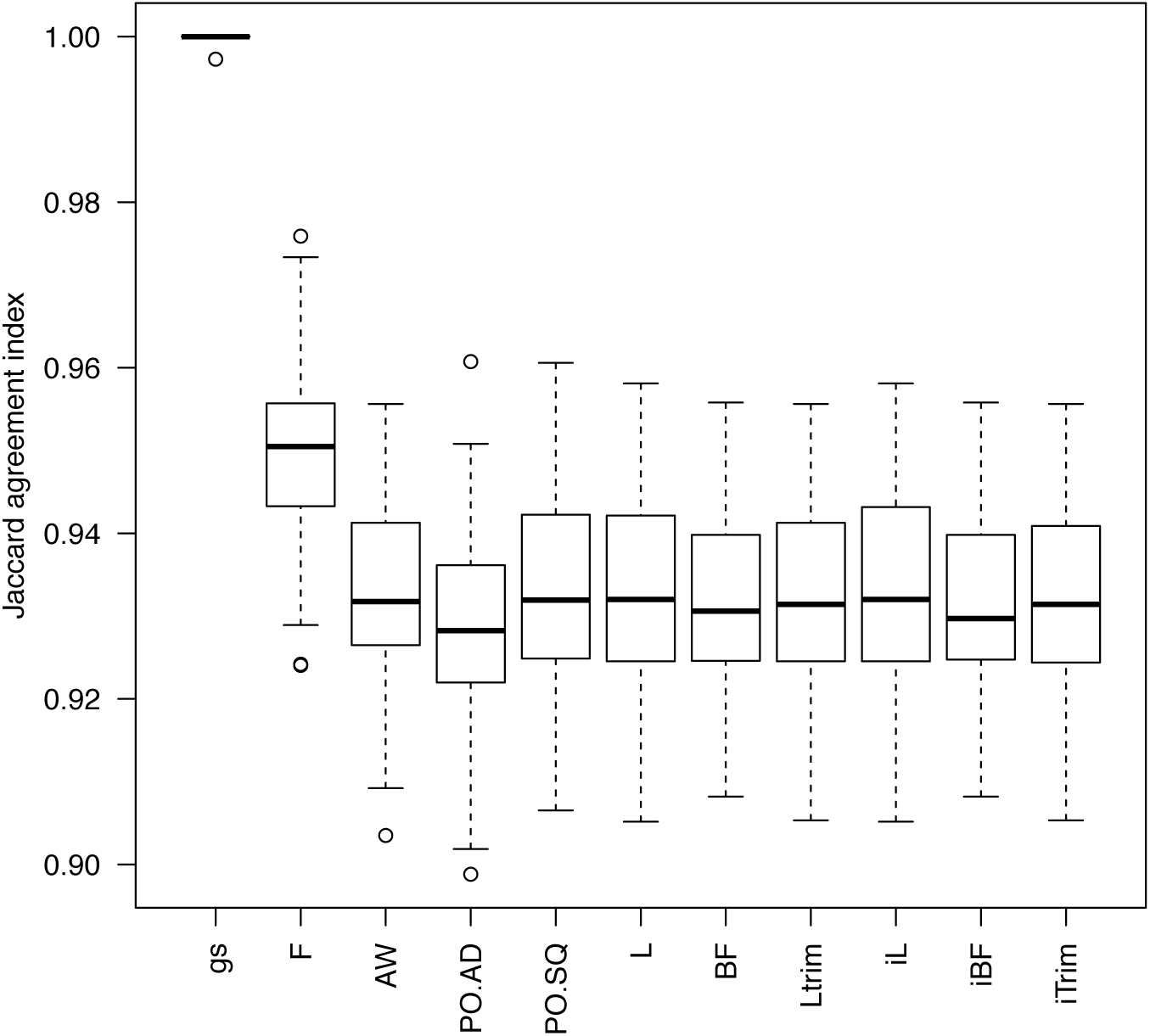
The boxplots of the 100 estimated Jaccard indices based on the 100 simulated datasets in SimI. The closer to one the Jaccrd index is, the better performance of a method

**Figure 3.**
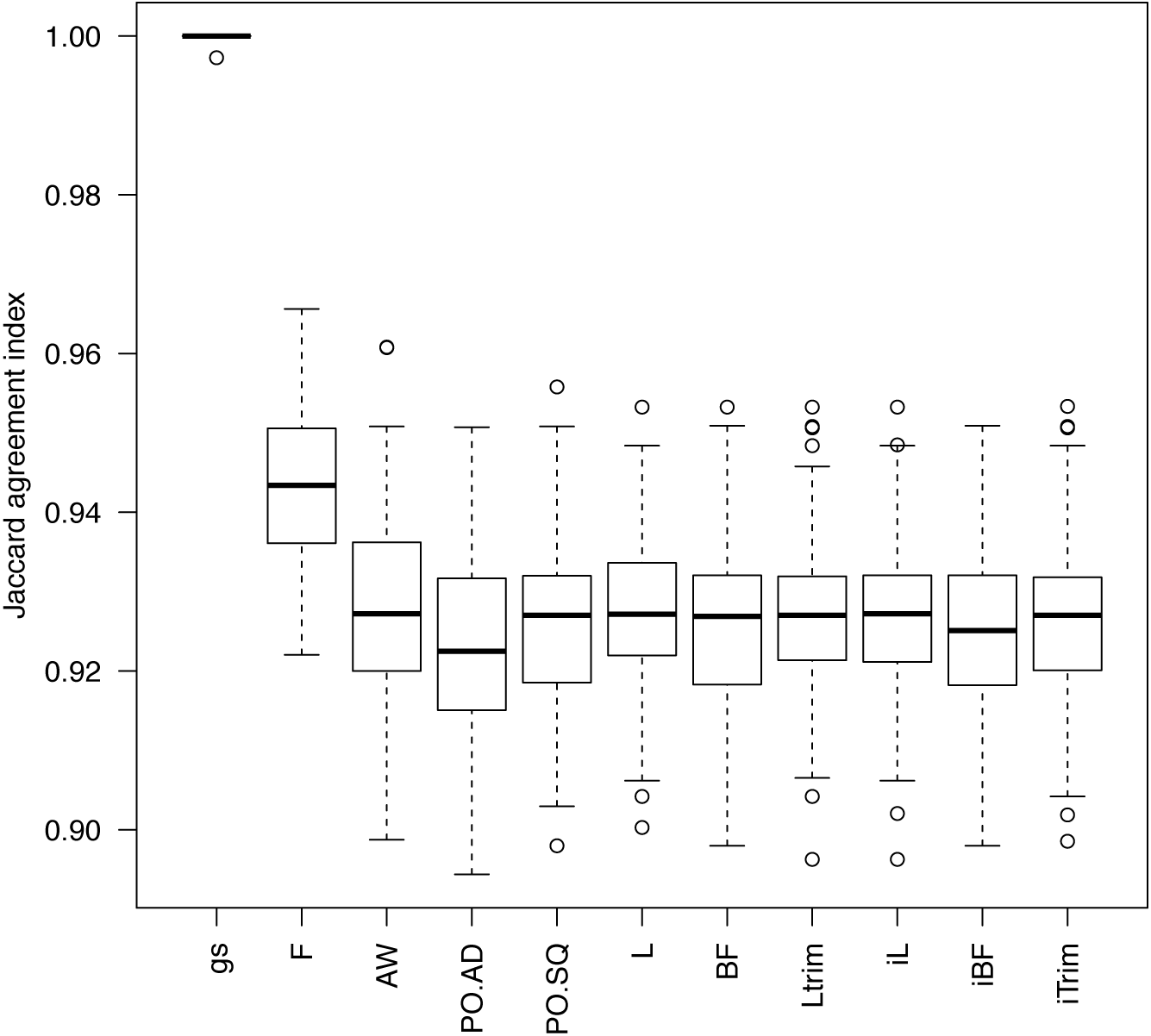
The boxplots of the 100 estimated Jaccard indices based on the 100 simulated datasets in SimIII. The cloer to one the Jaccard index is, the better performance of a method is.

The gs method still had much higher values of the Jaccard index than the 10 existing equal-variance tests even for data generated from multivariate t distributions, although the average values of Jaccard index were much smaller (<0.5) in SimII (Figure 2) and SimIV (Figure 4).

**Figure 2.**
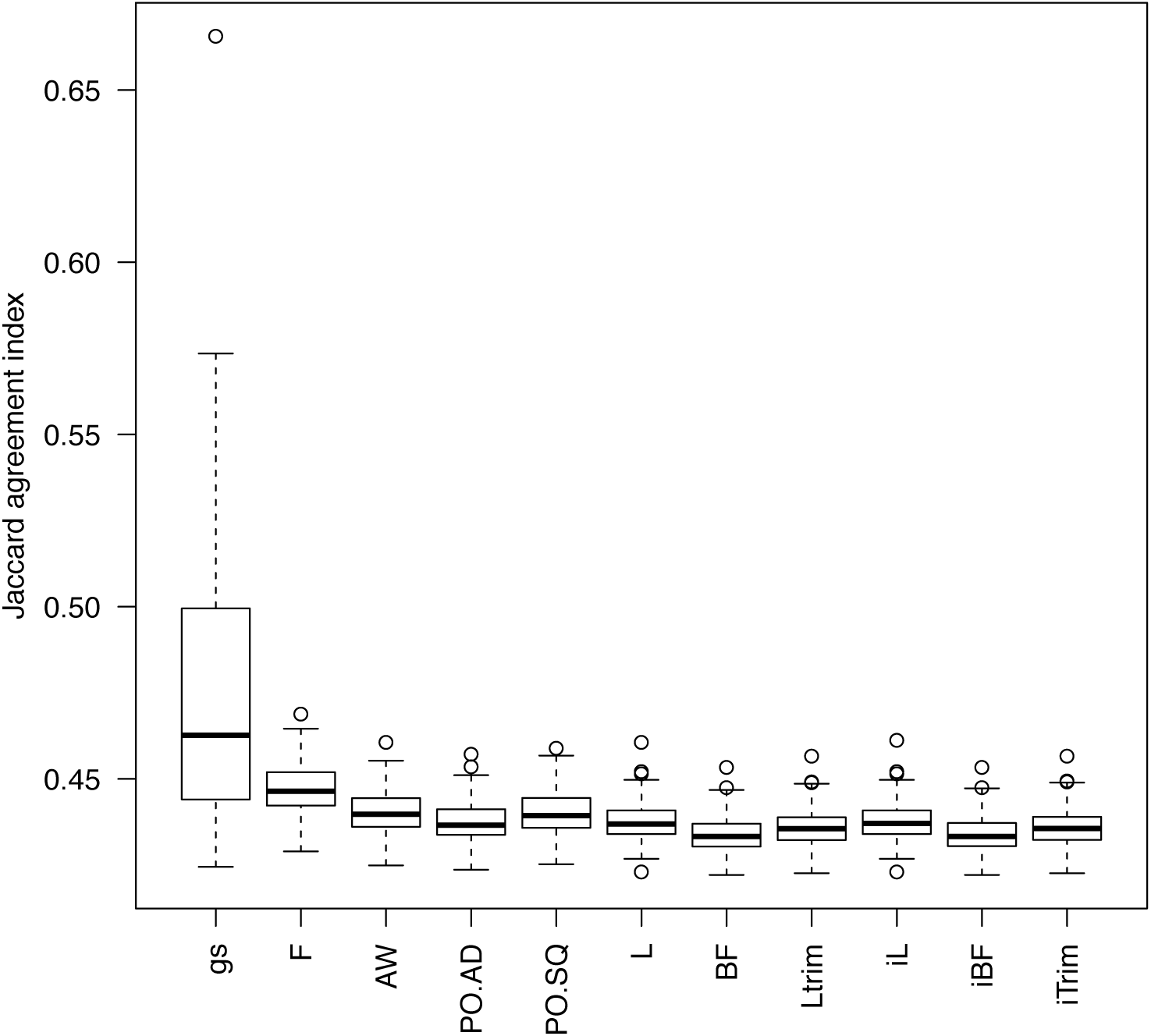
The boxplots of the 100 estimated Jaccard indices based on the 100 simulated datasets in SimII. The closer to one the Jaccard index is, the better performance of a method is.

**Figure 4.**
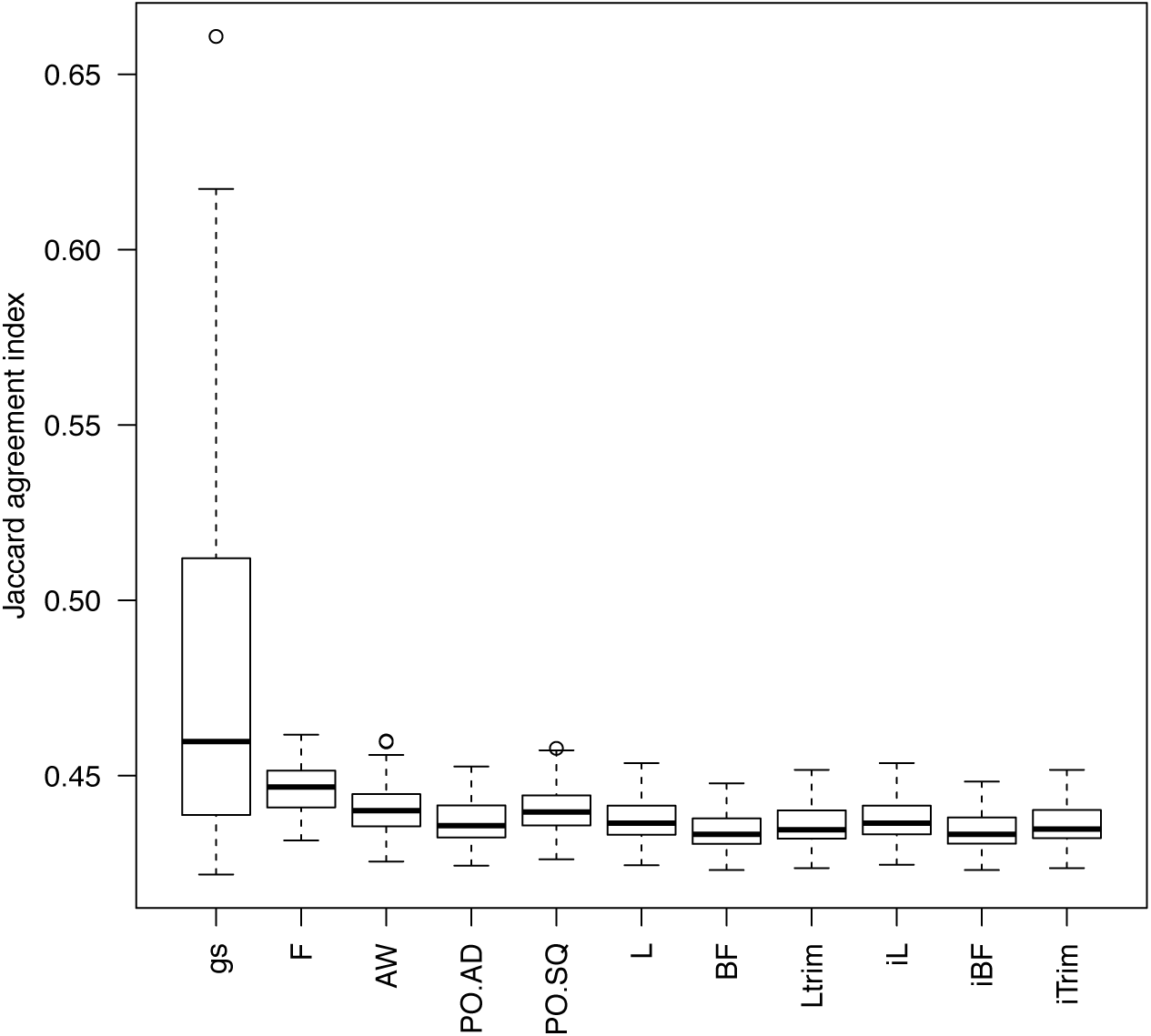
The boxplots of the 100 estimated Jaccard indices based on the 100 simulated datasets in SimIV. The closer to one the Jaccard index is, the better performance of a method is.

## Discussion

In this article, we proposed a novel method to detect microRNAs having different variances between cases and controls. The proposed method is different from probe-wised equal-variance tests in that it does not involve hypothesis testing since it is a model-based clustering. To the best of our knowledge, the proposed method is the first clustering algorithm to detect differential variable genomic probes.

In the 4 simulation studies, the proposed method outperformed 10 probe-wised tests, including the classic F test that has been reported to outperform other equal-variance tests when the normality assumption is held[17, 18]. The reason why the gs method performed better than the F test in SimI and SimIII, where the normality assumption is held, is that the gs method could borrow information across microRNAs (i.e., the estimation of the model parameters uses the information provided by al microRNAs).

The gs method and the F test outperformed the other 9 tests in SimII and SimIV, where the data were generated from multivariate t distribution (i.e., the normality assumption is violated). This indicates that the gs method and the F test could work well relative to other equal-variance tests for data with uni-modal and symmetric distributions. Fig. A6 shows the histograms and qqplots for a simulated dataset in each of the 4 simulation scenarios. In future research, we will evaluate the performance of the gs method in scenarios where the violation of the normality assumption is caused by skewed distributions. The robustness of the gs method warrants further investigation.

In the real data analysis, the gs method detected 67 validated DV microRNAs (66 OV and 1 UV), seven of which are DV only. The 7 DV-only microRNAs (hsa-miR-1826, hsa-miR-191, hsa-miR-194-star, hsa-miR-222, hsa-miR-502-3p, hsa-miR-93, and hsa-miR-99b) targeted to 1,639 genes based on the miRSystem analysis. Except for hsa-miR-1826, all DV-only microRNAs have been associated with HCC. Elyakim et al. (2010)[19] showed that miR-191 is a candidate oncogene target for hepatocellular carcinoma therapy. Law and Wong (2011)[20] reported the association of miR-194 with metastatic behavior of HCC. Murakami et al (2006)[21] reported that miR-222 is increased in poorly versus moderately versus well-differentiated hepatomas. Jin et al. (2016)[22] reported that miR-502-3p suppressed cell proliferation, migration, and invasion in HCC by targeting SET. Li et al. (2009)[23] confirmed that the miR-106b-25 cluster, which miR-93 belongs to, is over-expressed in HCC. Morishita et al. (2016)[24] found that miR-99b is up-regulated in HBV-infected HCC cells.

The 1,639 genes, which are targeted by the 7 DV-only microRNAs, are enriched in 6 KEGG pathways (CALCIUM SIGNALING PATHWAY, SALIVARY SECRETION, AMYOTROPHIC LATERAL SCLEROSIS (ALS), MAPK SIGNALING PATHWAY, PPAR SIGNALING PATHWAY, and ALZHEIMER’S DISEASE). All these 6 pathways have been linked to HCC in the literature. For example, Huang et al. (2017)[25] reported that increased mitochondrial fission induced cytosolic calcium signaling in HCC cells. Chen et al. (2017)[26] reported that in a mice study, DNA methylation marks that are differentially methylated between livers with HCC and livers without HCC are enriched in the SALIVARY SECRETION pathway. Seol et al.’s (2016)[27] results suggest that Riluzole, an amyotrophic lateral sclerosis (ALS) drug, has an anti-cancer effect on HCC. Feng et al. (2017)[28] reported that cantharidic acid inhibits HCC cell proliferation by inducing cell apoptosis through the p38 MAPK signaling pathway. Nwosu et al. (2017)[29] reported that down-regulated genes (HCC vs. non-HCC) were enriched in PPAR SIGNALING PATHWAY based on each of the 8 HCC datasets downloaded from the Gene Expression Omnibus (GEO). Jin et al. (2015)[30] reported that Kynurenine 3-monooxygenase (KMO), an enzyme playing a critical role in Huntington’s and Alzheimer’s diseases, exhibits tumor-promoting effects towards HCC. Hence, DV-only microRNAs are biologically relevant to HCC.

There are no overlaps among the enriched pathways for the 3 sets of microRNAs in the real data analysis: *S*_*DVonly*_ (the set of microRNAs that are validated DV, but not validated DE), *S*_*DEonly*_ (the set of microRNAs that are validated DE, but not validated DV), and *S*_*both*_ (the set of microRNAs that are both validated DE and validated DV). This indicates DV-only microRNAs might provide additional information about the molecular mechanisms of HCC than that provided by DE microRNAs.

Fig A7. showed that the distributions of the original real datasets are quite different from normal distributions. For the gs method, we followed Qiu et al.’s (2008) [6] data preprocessing. That is, we first performed the Box-Cox transformation, and then for each microRNA, we performed mean-centering and scaling operations so that the mean expression level is 0 and the variance is 1. Fig A7 showed that even after the Box-Cox transformation and scaling, the distributions of the data are still quite different from normal distributions. In the real data analysis, we applied the original data downloaded from GEO for all the 10 existing equal-variance tests. We also tried to apply the F test to the Box-Cox transformed data. The F test detected 159 DV microRNAs from the discovery set (GSE67318), but 472 DV microRNAs from the validation set (GSE67319), which is more than half of the 847 microRNAs. Further investigation is warranted.

In summary, the proposed gs method outperformed existing equal-variance tests in the simulation studies and detected biologically relevant microRNAs in a real data analysis of HCC data. The gs method is based on a mixture of multivariate normal distributions. Although the gs method showed some robustness to the violation of the normality assumption, in future we will study how to improve the gs method to make it robust to the violation of the normality assumption.

## Acknowledgement

The research is partially supported by the NSERC Discovery Grants and by the National Institute of Health of the United States (NIH/NHLBI 1 P01 HL 132825-01, NIH/NHLBI 5 R01 HL 125734)

## Supplementary Material

A concise description for each supplementary material file is shown in the table below:

| File | Description |
| --- | --- |
| suppl_document_MMDvariance3cat.pdf | Details about the gs model and its parameter estimation via the EM algorithm |
| TableA1.csv | Supplemental Table 1: The list of the 157 DE-only microRNAs |
| TableA2.csv | Supplemental Table 2: The list of the 60 DE-and-DE microRNAs |
| TableA3.csv | Supplemental Table 3: The list of the 1,639 genes targeted by the 7 microRNAs in $S_{DVonly}$ |
| TableA4.csv | Supplemental Table 4: The list of the 8,141 genes targeted by the 157 microRNAs in $S_{DEonly}$ |
| TableA5.csv | Supplemental Table 5: The list of the 6,893 genes targeted by the 60 microRNAs in $S_{both}$ |
| TableA6.csv | Supplemental Table 6: The list of the 6 KEGG pathways enriched by the 1,639 genes targeted by the 7 microRNAs in $S_{DVonly}$ |
| TableA7.csv | Supplemental Table 7: The list of the 1 KEGG pathway enriched by the 8,141 genes targeted by the 157 microRNAs in $S_{DEonly}$ |
| TableA8.csv | Supplemental Table 8: The list of the 2 KEGG pathways enriched by the 6,893 genes targeted by the 60 microRNAs in $S_{both}$ |
| FigsA1_A2.docx | A MS WORD file includes Fig A1 (Supplemental Figure 1) and Fig A2 (Supplemental Figure 2).<br>Fig A1. The plot of percentiles of log2 expression levels across arrays. Left panel: GSE67138; Right panel: GSE67139. |
|  | Fig A2. The plot of the first and second principal components. Left panel: GSE67138; Right panel: GSE67139. |
| FigA3.pdf | Supplemental Figure 3: Parallel boxplots of the 7 validated DV-only microRNAs. Left panel: GSE67138; Right panel: GSE67139 |
| FigA4.pdf | Supplemental Figure 4: Parallel boxplots of the 157 validated DE-only microRNAs. Left panel: GSE67138; Right panel: GSE67139 |
| FigA5.pdf | Supplemental Figure 5: Parallel boxplots of the 60 validated DV-and-DE microRNAs. Left panel: GSE67138; Right panel: GSE67139 |
| FigA6.pdf | Supplemental Figure 6: Histograms and QQ plots for a simulated dataset in each of the 4 simulation scenarios |
| FigA7.pdf | Supplemental Figure 7: Histograms and QQ plots for the two real datasets |

## Reference

1. Hansen, K.D., et al., Increased methylation variation in epigenetic domains across cancer types. Nat Genet, 2011. 43(8): p. 768–75.

2. Jaffe, A.E., et al., Significance analysis and statistical dissection of variably methylated regions. Biostatistics, 2012. 13(1): p. 166–78.

3. Teschendorff, A.E. and M. Widschwendter, Differential variability improves the identification of cancer risk markers in DNA methylation studies profiling precursor cancer lesions. Bioinformatics, 2012. 28(11): p. 1487–94.

4. Hayes, J., P.P. Peruzzi, and S. Lawler, MicroRNAs in cancer: biomarkers, functions and therapy. Trends Mol Med, 2014. 20(8): p. 460–9.

5. Simonson, B. and S. Das, MicroRNA Therapeutics: the Next Magic Bullet? Mini Rev Med Chem, 2015. 15(6): p. 467–74.

6. Qiu, W., et al., A marginal mixture model for selecting differentially expressed genes across two types of tissue samples. Int J Biostat, 2008. 4(1): p. Article 20.

7. Dempster, A., N. Laird, and D. Rubin, Maximum likelihood from incomplete data via the EM algorithm. Journal of the Royal Statistical Society, Series B, 1977. 39: p. 1–38.

8. Edgar, R., M. Domrachev, and A.E. Lash, Gene Expression Omnibus: NCBI gene expression and hybridization array data repository. Nucleic Acids Res, 2002. 30(1): p. 207–10.

9. Ahn, S. and T. Wang, A powerful statistical method for identifying differentially methylated markers in complex diseases. Pac Symp Biocomput, 2013: p. 69–79.

10. Phipson, B. and A. Oshlack, DiffVar: a new method for detecting differential variability with application to methylation in cancer and aging. Genome Biol, 2014. 15(9): p. 465.

11. Levene, H., Robust tests for equality of variances, in Contributions to Probability and Statistics: Essays in Honor of Harold Hotelling, O. I., Editor. 1960, Stanford University Press. p. 278–292.

12. Brown, M.B. and A.B. Forsythe, Robust tests for the equality of variances. Journal of the American Statistical Association, 1974. 69: p. 364–367.

13. Qiu, W., et al., New Score Tests for Equality of Variances in the Application of DNA Methylation Data Analysis. Insights in Genetics and Genomics, 2016. 1: p. 3.1.

14. Ritchie, M.E., et al., limma powers differential expression analyses for RNA-sequencing and microarray studies. Nucleic Acids Res, 2015. 43(7): p. e47.

15. Lu, T.P., et al., miRSystem: an integrated system for characterizing enriched functions and pathways of microRNA targets. PLoS One, 2012. 7(8): p. e42390.

16. Jaccard, P., The distribution of the flora in the alpine zone. New Phytologist, 1912. 11: p. 37–50.

17. Conover, W.J., M.E. Johnson, and M.M. JohnSon, A Comparative Study of Tests for Homogeneity of Variances, with Applications to the Outer Continental Shelf Bidding Data. Technometrics, 1981. 23(4): p. 351–361.

18. Li, X., et al., A Comparative Study of Tests for Homogeneity of Variances with Application to DNA Methylation Data. PLoS One, 2015. 10(12): p. e0145295.

19. Elyakim, E., et al., hsa-miR-191 is a candidate oncogene target for hepatocellular carcinoma therapy. Cancer Res, 2010. 70(20): p. 8077–87.

20. Law, P.T. and N. Wong, Emerging roles of microRNA in the intracellular signaling networks of hepatocellular carcinoma. J Gastroenterol Hepatol, 2011. 26(3): p. 437–49.

21. Murakami, Y., et al., Comprehensive analysis of microRNA expression patterns in hepatocellular carcinoma and non-tumorous tissues. Oncogene, 2006. 25(17): p. 2537–45.

22. Jin, H., et al., MiR-502-3P suppresses cell proliferation, migration, and invasion in hepatocellular carcinoma by targeting SET. Onco Targets Ther, 2016. 9: p. 3281–9.

23. Li, Y., et al., Role of the miR-106b-25 microRNA cluster in hepatocellular carcinoma. Cancer Sci, 2009. 100(7): p. 1234–42.

24. Morishita, A., et al., MicroRNA profiles in various hepatocellular carcinoma cell lines. Oncol Lett, 2016. 12(3): p. 1687–1692.

25. Huang, Q., et al., Mitochondrial fission forms a positive feedback loop with cytosolic calcium signaling pathway to promote autophagy in hepatocellular carcinoma cells. Cancer Lett, 2017. 403: p. 108–118.

26. Chen, H., et al., Hepatic cyclooxygenase-2 overexpression induced spontaneous hepatocellular carcinoma formation in mice. Oncogene, 2017. 36(31): p. 4415–4426.

27. Seol, H.S., et al., Glutamate release inhibitor, Riluzole, inhibited proliferation of human hepatocellular carcinoma cells by elevated ROS production. Cancer Lett, 2016. 382(2): p. 157–165.

28. Feng, I.C., et al., Cantharidic acid induces apoptosis through the p38 MAPK signaling pathway in human hepatocellular carcinoma. Environ Toxicol, 2017.

29. Nwosu, Z.C., et al., Identification of the Consistently Altered Metabolic Targets in Human Hepatocellular Carcinoma. Cell Mol Gastroenterol Hepatol, 2017. 4(2): p. 303–323.e1.

30. Jin, H., et al., Prognostic significance of kynurenine 3-monooxygenase and effects on proliferation, migration, and invasion of human hepatocellular carcinoma. Sci Rep, 2015. 5: p. 10466.

